# Removal of the extracellular matrix biases auditory cortical layer dynamics towards supragranular frequency integration

**DOI:** 10.1101/2020.05.04.076224

**Authors:** M El-Tabbal, H Niekisch, JU Henschke, E Budinger, R Frischknecht, M Deliano, MFK Happel

## Abstract

In the adult vertebrate brain, enzymatic removal of the extracellular matrix (ECM) is increasingly recognized to promote learning, memory recall, and restorative plasticity. However, the underlying impact of local removal of the ECM on cortical circuit processing is still not understood. Here, we removed the ECM in the primary auditory cortex (ACx) of adult Mongolian gerbils using local injections of hyaluronidase (HYase). Using laminar current-source density (CSD) analysis, we found layer-specific changes of the spatiotemporal synaptic activity patterns. Applying multitaper spectral analysis and time-domain conditional Granger causality (GC) measures, we found increased stimulus-evoked oscillatory power in the beta band (25-36 Hz) selectively within infragranular layers Vb and enhanced supragranular synaptic activity. Our findings reveal new insights on how ECM modulation affects the sensory integration via altered translaminar cortical network dynamics with a supragranular lead of the columnar response profile.

## Introduction

While basic neuronal networks established during development have to be conserved, their activity-dependent fine-tuning and modification form the basic mechanism for adult learning and memory. In the vertebrate brain, long-lasting structural tenacity of neuronal networks is supported by the extracellular matrix (ECM). It consists of the core backbone glycan hyaluronic acid that attaches chondroitin sulfate proteoglycans and other proteins (Fawcett et al., 2019a; Frischknecht and Seidenbecher, 2012). The most prominent forms of the adult ECM in the brain are *i)* perineuronal nets (PNN) named by their mesh-like appearance around cell bodies and proximal dendrites of mainly parvalbumin-positive (PV+) interneurons, and *ii)* the “loose” ECM found almost ubiquitously in the brain (Fawcett et al., 2019a; Patton et al., 2019). The ECM is formed during brain maturation implementing a switch from juvenile to adult forms of synaptic plasticity (Gundelfinger et al., 2010; Pizzorusso et al., 2002). For instance, enriched PNNs have been linked to states of reduced structural plasticity (de Vivo et al., 2013; Dick et al., 2013) In a seminal study, Pizzorusso and colleagues demonstrated that enzymatic removal using chondroitinase ABC reinstalls critical period plasticity (Pizzorusso et al., 2002). ECM removal within the cortex or hippocampus has further diverse effects on contextual fear conditioning (Banerjee et al., 2017; Hylin et al., 2013), object recognition (Romberg et al., 2013), extinction of fear or drug memories (Gogolla et al., 2009; Xue et al., 2014), and auditory reversal learning (Happel et al., 2014b). During learning processes in the developing and adult brain, variability of ECM densities and dynamic remodeling has been shown to support learning-dependent plasticity (Cornez et al., 2020; Niekisch et al., 2019). In accordance, systemic injection of dopamine receptor D1 agonists *in vivo* promote rapid cleavage of the ECM protein brevican (Mitlöhner et al., 2020). ECM weakening increases lateral diffusion of glutamate receptors at the synapse (Frischknecht et al., 2009; Heine et al., 2008; Pandya et al., 2018; Schweitzer et al., 2017) and alters the balance between inhibition and excitation (Lensjø et al., 2017). However, the mechanistic understanding on how states of reduced ECM may affect circuit processing in a layered structure, such as the cortex, is rather elusive.

Here, we acutely weakened the ECM in the primary auditory cortex (ACx) of Mongolian gerbils (*Meriones unguiculatus*) by local microinjections of hyaluronidase (HYase). In parallel, we used laminar recordings of local field potentials and current-source density (CSD) analysis (Happel et al., 2010; Happel and Ohl, 2017) to quantify the spatiotemporal sequence of spontaneous and stimulus-evoked laminar synaptic activation. Local HYase injection led to generally stronger tone-evoked synaptic currents in supragranular layers I/II and a stimulus-specific decrease in infragranular layer Vb. Synaptic activity was hence imbalanced between these two layers, with a general stimulus-independent activation shift towards supragranular layers I/II, which was accompanied by an increase of lateral corticocortical input revealed by CSD residual analysis (Happel et al., 2010). Utilizing a multitaper spectral analysis of tone-evoked oscillatory circuit responses (Deliano et al., 2018), we revealed a layer specific change of the oscillatory power in the beta band (25-36 Hz) within infragranular layers Vb. To investigate how these changes are mediated by the translaminar dynamics of cortical network activity, we implemented time-domain Granger causality (GC) measures mapping the directional causal relationship between activity across cortical layers. After application of HYase, we found a significant increase in the GC from supragranular layers I/II onto infragranular layer Vb. Additionally, there was a significant decrease in the infragranular layer VI drive routing back towards granular input layers III/IV. Our data thereby shows that enzymatic removal of the ECM acutely biases the columnar synaptic network processing towards stronger recruitment of supragranular circuits and enhancement of lateral, crosscolumnar interactions and uncoupling of deeper layer circuits. Our study provides a mesocopic cortical circuit mechanism of enhanced sensory integration in upper layers, that might help to better understand existing behavioral findings linked to ECM modulation and may further allow to optimize concepts for therapeutic approaches targeting the ECM.

## Results

In the present study, the extracellular matrix was locally removed in unilateral primary auditory cortex of adult Mongolian gerbils using microinjection of the ECM-degrading enzyme hyaluronidase. By placing a linear multichannel recording electrode in the near proximity (<250 μm) of the glass capillary, we recorded the laminar local field potential across all layers within the region of the weakened ECM. CSD analysis was used to investigate how ECM removal acutely affects laminar tone-evoked processing and spontaneous activity. Specifically, analysis of the relative residual of the CSD, layer-specific multitaper spectral analysis and Granger causality measures allowed us to identify enhanced corticocortical input in upper layers to dominate the translaminar processing dynamics within the cortical column (Happel et al., 2010; Happel and Ohl, 2017).

### Layer-dependent changes of tone-evoked synaptic activity

Pure tone-evoked laminar CSD profiles were first recorded in the untreated primary auditory cortex. Recordings showed a canonical feedforward processing pattern with main short-latent current sink components in granular layers III/IV and infragranular layer Vb most prominently for stimulation with the BF (Fig. 1A). Those short-latent sinks reflect the synaptic afferent input from the vMGB with main projections to granular layers and collaterals within layer Vb (Brunk et al., 2019; Happel et al., 2014a; Schaefer et al., 2015). Early sinks were followed by synaptic activation of supragranular (I/II) and deep infragranular (VI) layers (Fig. 1) which are due to intracortical synaptic processes (Happel et al., 2010). Then, the ECM-degrading enzyme HYase was injected in close proximity at the recording site (the micropipette was inserted ~250μm next to the electrode). Before the start of the recording and tissue was allowed to recover from insertion for >1h. After HYase injection we again waited for >2.5h in order to let the enzyme effectuate the ECM in the proximity of the recording patch. In a previous study, we showed that by local injection at one location the enzyme effectuates a region with a diameter of around 1mm well covering the recording site (Happel et al., 2014b). After ECM removal, the feedforward CSD pattern changed in a layer-dependent manner. Tone evoked synaptic responses showed a decrease of early infragranular input and a parallel increase of supragranular current flow (Fig. 1). No obvious differences were found for layers III/IV and VI. We stained brains after local unilateral HYase injection in temporal cortex against mouse anti-PV (green) and anti-WFA (red) staining (Fig. 1). High-resolution confocal microscopy (Zeiss LSM 700, Germany) revealed a layer-dependent accumulation of PNN structures in the ACx particularly around PV+ interneurons in cortical layers III/IV of the control injection side, which were effectively reduced at the side of HYase injection. We further quantified tone-evoked RMS amplitudes of individual cortical layer activity as a function of stimulation frequency (Fig. 2A). Pure-tone derived frequency response curves showed a consistent decrease of sink activity in infragranular layer Vb induced by HYase injection. The effect was stronger at BF stimulation compared to off-BF stimulation, which indicates a reduced sharpness of spectral tuning. A 2-way rmANOVA showed significant main effects for factors tone-frequency (‘Freq’), HYase treatment (‘HYase’) and a significant interaction ‘Freq x HYase’ (see Tab. 1). Thus, besides an overall reduction of the Vb response by HYase as main-effect, there was a frequency-specific interaction effect that cannot be explained only by a general modulation of overall excitability. In contrast, layer I/II RMS-amplitude was increased in a frequency-specific manner (2-way rmANOVA with main effect for ‘Freq’ and a significant interaction ‘Freq x HYase’; Tab. 1). Granular layer III/IV and infragranular VI activity was unchanged after HYase injection.

**Fig. 1.**
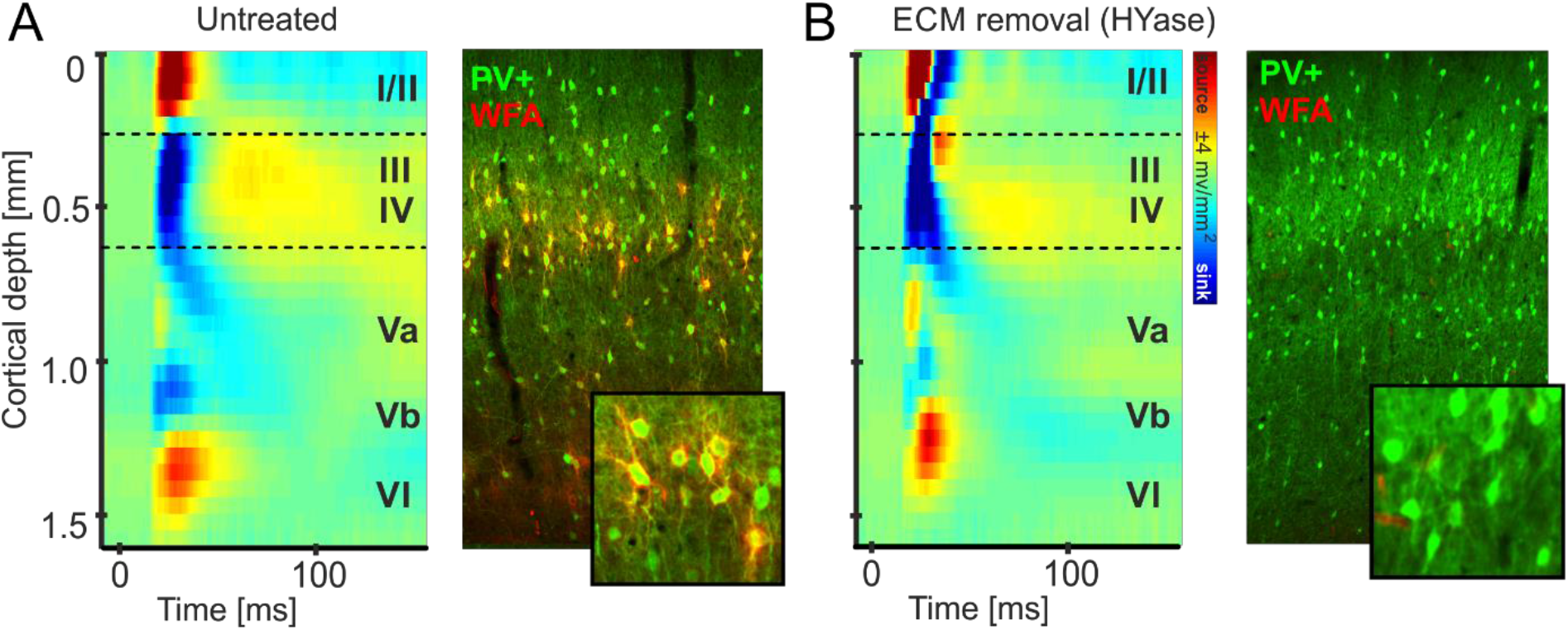
ECM weakening changed tone-evoked columnar processing in a layer-dependent manner. **A.** Tone-evoked CSD profiles (left) display a canonical feedforward pattern with afferent early sink activity in granular and infragranular layer Vb and subsequent activation of supragranular and infragranular layers. Roman numbers indicate cortical layers. High-resolution confocal microscopy images (Zeiss LSM 700, Germany) show mouse anti-PV (green) and anti-*WFA* (red) staining in the vicinity of the recording site (right). Stainings reveal layer-dependent accumulation of PNN structures in the auditory cortex (see inset for details). **B.** After ECM removal by microinjections of HYase, tone-evoked CSD profiles showed reduced strength of the early afferent input in layer Vb and stronger activation of supragranular layers I/II. Immunostainings after HYase treatment also revealed reduced density of *WFA* and hence reduced PNNs around PV+ interneurons.

**Fig. 2.**
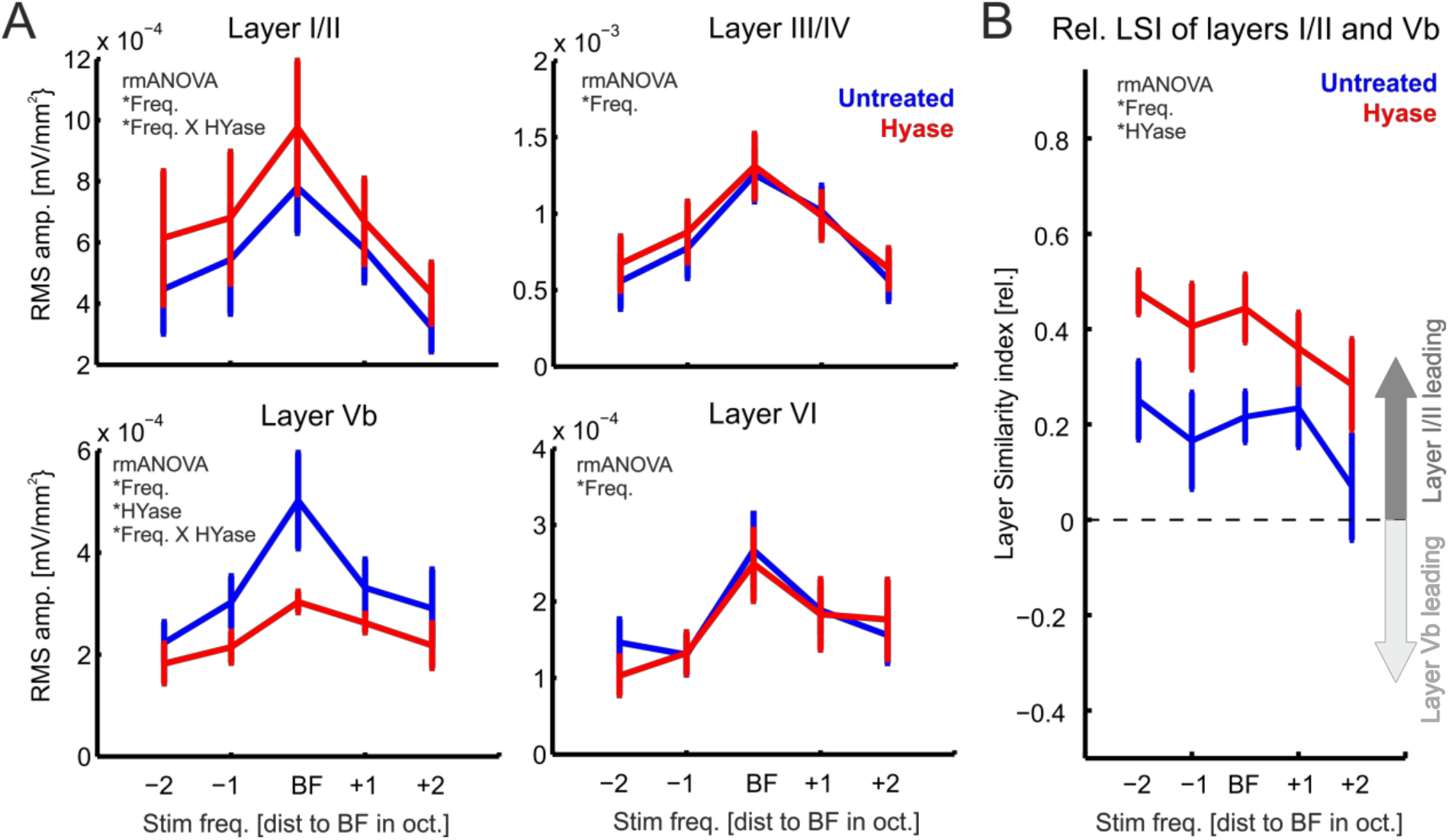
Removal of the ECM affects layer-specific processing in the auditory cortex. **A.** Frequency-response tuning curves for the RMS amplitude (±SEM) of CSD traces from individual cortical layers showed increased activity within supragranular layers I/II and decreased activity in cortical layer Vb. Activity in layers III/IV and VI showed did not change. **B.** Correspondingly, the Layer-Symmetry-Index LSI = (I/II – Vb) / (I/II + Vb) showed that after HYase treatment evoked synaptic activity shifted towards supragranular layers independent of the stimulation frequency. Significant main effects and interactions of a 2-way-rmANOVA are indicated in each subpanel by the asterisk * (for results see Tab. 1).

**Tab. 1.**
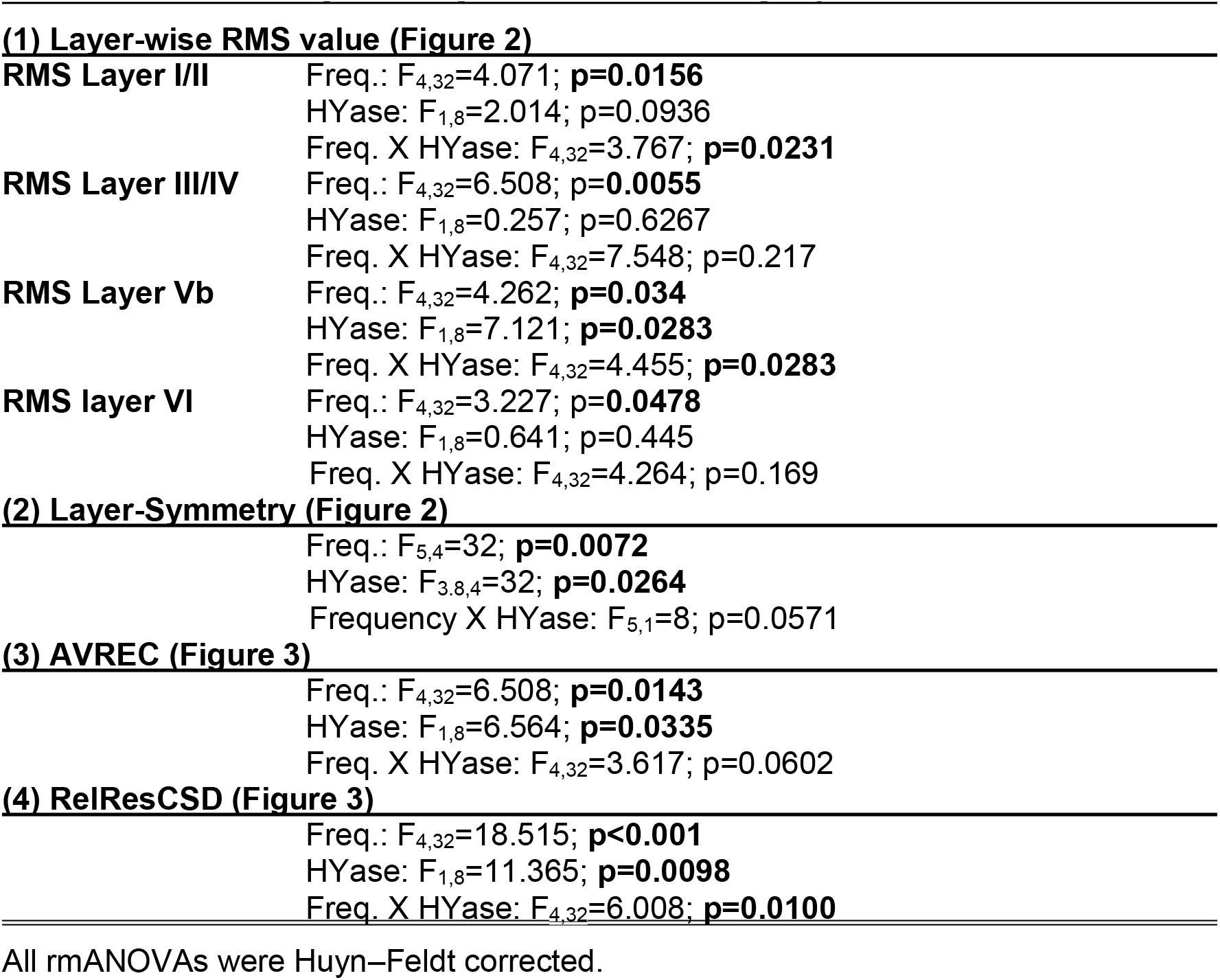
Statistical analysis of layer-wise CSD data by repeated-measures ANOVA.

To relate the observed changes in layer Vb and I/II to each other we calculated and quantified the relative shift of translaminar columnar activity, we calculated a Layer-Symmetry-index (LSI) of current flow between cortical layers I/II and Vb (cf. Materials and Methods;). By calculating the LSI = (I/II – Vb) / (I/II + Vb), positive LSI indicate stronger activation of supragranular layers compared to infragranular layers. In untreated cortex, supragranular layers generally display larger evoked synaptic currents compared to infragranular layers with LSI-value around +0.2 (Happel et al., 2014a). HYase treatment, however, amplified this supragranular lead of current flow and LSI values exceeded values of +0.4. Correspondingly, a 2-way rmANOVA of LSI of RMS amplitudes showed main effects for ‘Freq’ and ‘HYase’ without interaction (Tab. 1). This shows, that the frequency specific decrease of responses in layer Vb and increase in layer I/II were counterbalanced, and that the response ratio increased across all frequencies.

Supragranular sink activity has been related to horizontal, intercolumnar processing (Happel et al., 2014; Kaur et al., 2004). In order to further quantify crosscolumnar activity spread, we analyzed the frequency-response tuning of the AVREC and the RelResCSD (Fig. 3). The AVREC showed a moderate, but consistent increase after HYase injection (2-way rmANOVA with main effects ‘Freq’ and ‘HYase’). The RelResCSD showed a likewise increase that dependent on frequency revealed by a 2-way rmANOVA with main effect ‘Freq’ and a significant interaction of ‘Freq x Treatment’ (see Tab. 1). Both findings are in accordance with increased supragranular activity as a relative indicator for synaptic input from neighboring cortical columns. Correspondingly, the Q40dB response bandwidth of the AVREC did not change significantly, while the response bandwidth of the RelResCSD was significantly increased. Hence, the higher overall activity measured by the AVREC was mainly due to a stronger activation found in supragranular layers I/II leading to an increased corticocortical activity spread measured by the RelResCSD. Comparable changes of the columnar response profile were not found after injection of 0.9% sodium chloride in a set of control animals (n=3; Suppl Fig. 1).

**Fig. 3.**
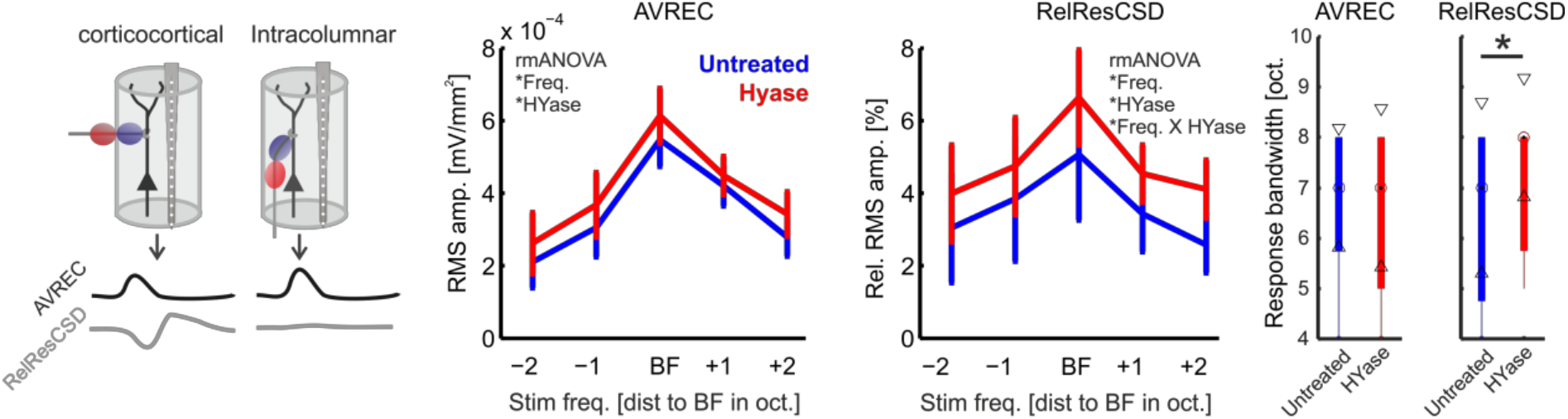
Quantification of AVREC and RelResCSD frequency tuning after ECM removal. *Left,* Schema illustrating that AVREC activity is mainly associated with local columnar activity, while the RelResCSD quantifies contributions from lateral corticocortical input (cf. Happel et al., 2014, 2010). Frequency-response tuning curves of AVREC RMS amplitudes (±SEM) showed increased activity across all stimulation frequencies. RelResCSD showed increase that was depending on stimulation frequency. *Right*, Correspondingly, AVREC showed no change of Q40dB bandwidth, while RelResCSD bandwidths were significantly increased (*paired Student’s t-test, p=0.02). Significant main effects and interactions of a 2-way-rmANOVA are indicated in each subpanel by the asterisk * (for results see Tab. 1).

### Spectral power of tone-evoked processing changed mainly in infragranular layers

Next we were interested in the underlying oscillatory nature of the layer-specific synaptic activity. We found that ECM removal had a layer-dependent impact on the spectral power distribution within exclusively infragranular layer Vb. Fig. 4A shows the grand mean power spectrum (±SEM) of the averaged individual cortical layer activity after BF stimulation (evoked spectrum). To avoid the bias in selecting frequency bands we used the t-values statistical comparison to detect the spectral difference between the experimental conditions. HYase treatment caused significant increase in the evoked spectral responses in infragranular layer Vb in the beta range from 25–36 Hz. Spectral power effects might include background effects unrelated to the stimulus presentation. We therefore further calculated the power spectrum during spontaneous recordings. Fig. 4 shows the t-values calculated as the mean spectral differences between these conditions across subjects (n = 9) and tapers (nt = 5) normalized by their standard error (SEM). The spectrum of spontaneous activity showed no significant difference before and after HYase injection.

**Fig. 4.**
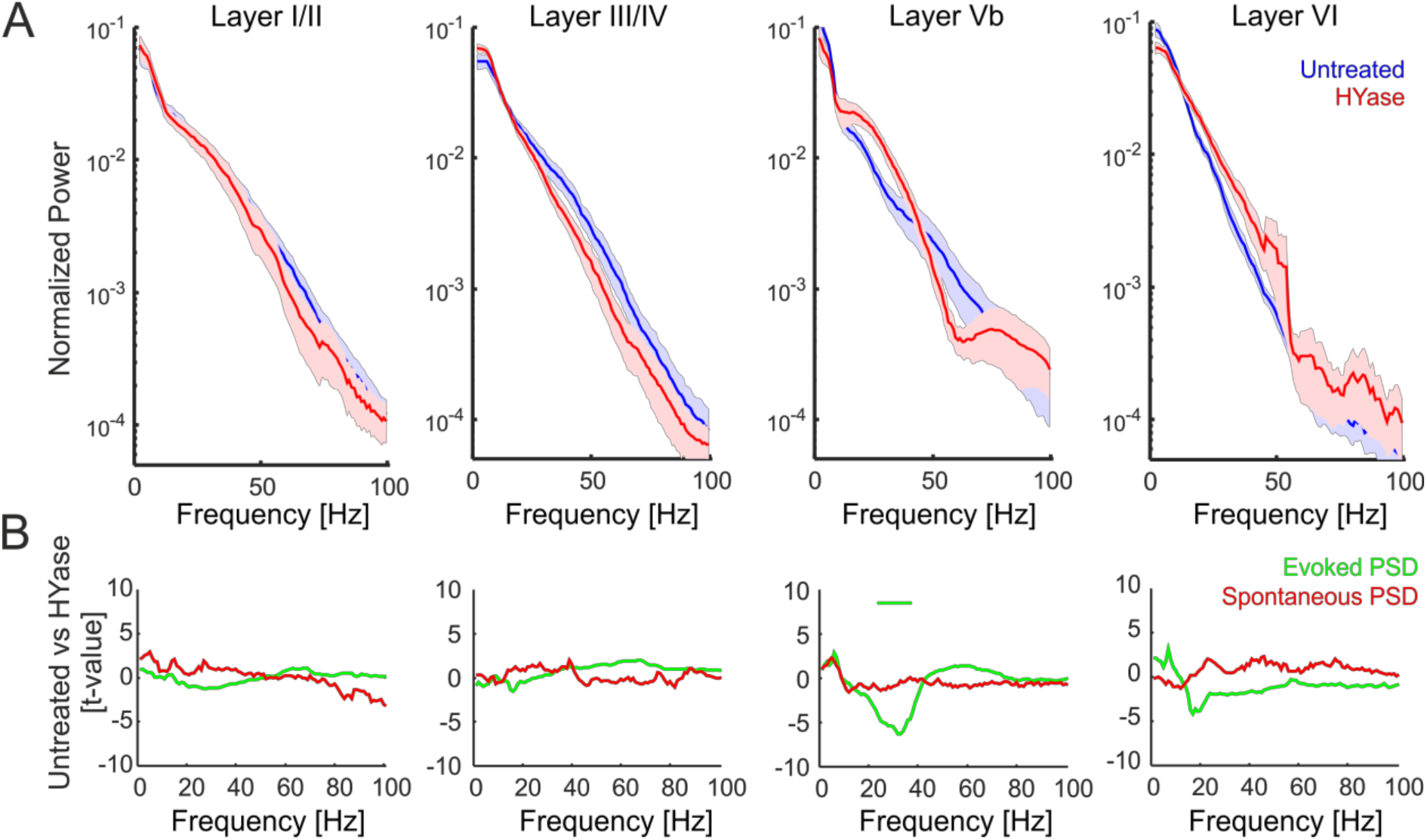
Analysis of alterations in layer-dependent spectral power after HYase treatment. **A.** Layer-dependent analysis of frequency components of evoked responses showed bimodal changes in the spectral power distribution mainly in infragranular layers. In both early and late infragranular layers, beta power showed a significant increase. **B.** Standardized differences of evoked (green) and spontaneous (red) power spectra between untreated recordings and after HYase treatment expressed as t-values for each frequency bin (negative t-values indicate higher power after HYase injection). Power spectra were log-transformed for each taper on the single-trial level. Standardization was based on sample means and standard errors of the mean across subjects (n = 8) and tapers (nt = 5). Asterisks mark the significant differences for the beta range from 25-36 Hz for the evoked PSD in infragranular layer Vb by a point-by-point Students-t-test comparison with a Benjamin Hochberg correction controlling for false discovery rates.

### Response dynamics across cortical layers

We hypothesized that the temporal relationship between activity measured in cortical layers was changed after application of HYase. We further assume that the columnar response was modulated by a differential impact of supragranular and infragranular layers on the laminar response dynamics. To test this, we analyzed this dynamical relationship by employing time-domain Granger causality (GC) estimation on the single trial CSD traces from each cortical layer to generate a network model for each animal and then compared the granger causality matrices. Fig. 5A depicts the directed pairwise G-causal estimates between cortical layers by the thickness of connecting arrows. Fig. 5B shows the corresponding mean values of the G-causal estimate. In order to compare laminar dynamics before and after application of HYase, we compared the pairwise conditional GC measures. We found a significantly increased predictive power of supragranular activity for activity recorded in infragranular layer Vb. Further, predictive power of infragranular layer VI activity for granular layer III/IV activity was significantly reduced during evoked columnar responses. This was in contrast to the significant strengthening of drive of infragranular layer Vb to the infragranular layer VI (Fig. 5C).

**Fig. 5.**
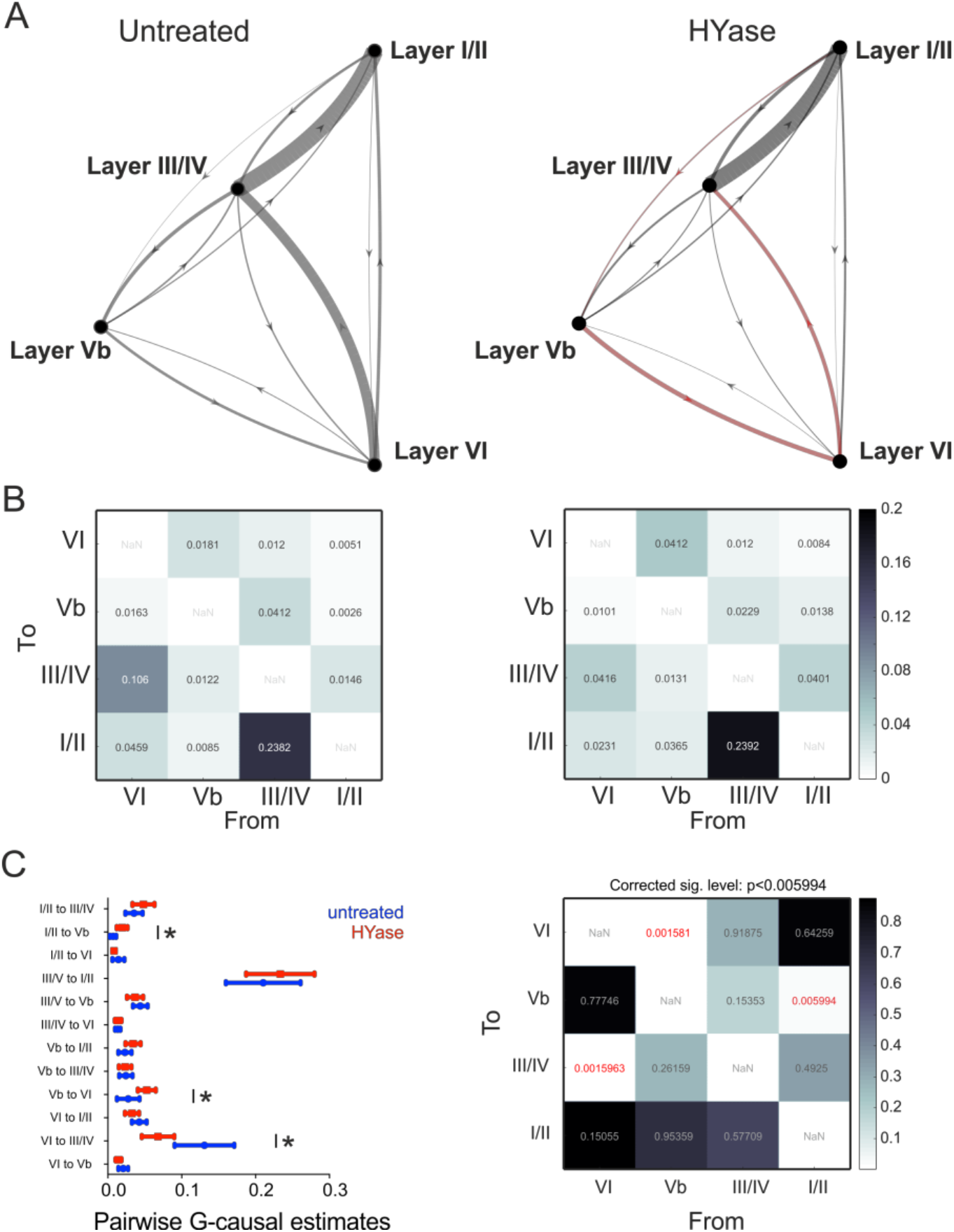
Analysis of time-domain pairwise-conditional granger causality. Granger causality matrices were calculated based on the single trial CSD traces from each cortical layer for each subject (n=9). **A.** Mean values of the granger causality estimates are shown before (*left*) and after (*right*) HYase treatment. The dynamics of the predictive causal relationship in different cortical layers before and after HYase injection is depicted as a G-causal network graph. Each node represents a single cortical layer and the arrows are the directed causal relation between different layers, with arrow thickness indicating the mean of the causal estimate weights across animals. The red arrows in the right GC plot indicate where G-causal forecasting undergoes significant changes after HYase treatment. **B**. The corresponding mean values of the conditional pairwise G-causal matrix across animals before (*left*) and after (*right*) HYase treatment. **C.** Comparison of the time-domain pairwise-conditional granger causality in the two conditions before and after HYase injection. *Left*. G-causal estimates were plotted with mean and standard error of the mean for each layer pairs from the 9 animals. *Right*, A multiple paired t-test was done on the log transformed values of the estimates and then corrected based on a false discovery rate of 0.05. Significant differences between G-causal measures indicated in the left plot correspond to a corrected level of significance after multiple comparisons of p<0.003558.

### Layer-dependent changes of spontaneous columnar activity

Finally, we wanted to investigate, whether the aforementioned changes of the columnar responses are specific for sensory evoked activity. We therefore recorded periods of 6 s of spontaneous activity in individual animals (n=7) before and after HYase-induced removal of the ECM. CSD recordings in the untreated auditory cortex revealed the occurrence of spontaneous events of translaminar synaptic information flow, which we refer to as spontaneous columnar events (SCE). SCE’s showed a comparable, but distinct synaptic current flow compared to tone-evoked columnar feedforward activation patterns. While tone-evoked responses display earliest synaptic inputs in both granular layers IV and infragranular layer Vb due to afferent thalamocortical input, SCE’s showed initial synaptic activity in layer Vb initiating translaminar information flow (Sakata and Harris, 2009). Activity in layer Vb was followed by current sink translaminar synaptic activity across the entire column including infragranular and supragranular layers (Fig. 6A, *top*). For quantification of the frequency of SCE’s we used a peak detection analysis based on single-trial AVREC traces (see Methods; Fig. 6A, *bottom*). After microinjection of HYase in the vicinity of the recording axis, spontaneous CSD profiles changed considerably. We still observed SCE’s, however, with an altered current flow of columnar activity. Individual SCE’s were shorter in duration and showed significantly reduced synaptic current flow across all layers (Fig. 6B). ECM weakening also significantly reduced the rate of SCE/s (Fig. 6C). We further attempted to quantify the overall cortical activation based on the overall current flow within the column quantified by the RMS amplitude of the AVREC. We compared this with the RelResCSD quantifying the unbalanced contributions of sinks and sources of the CSD profile, as an indicator for corticocortical spread of activity across columns (see Fig. 3; cf. Happel et al., 2010). While the AVREC showed a consistent significant reduction after HYase injection, the RelResCSD showed a parallel increase indicating relative higher crosscolumnar activity spread after treatment (Fig. 6D), which is in accordance with the tone-evoked activity reported before (Fig. 3).

**Fig. 6.**
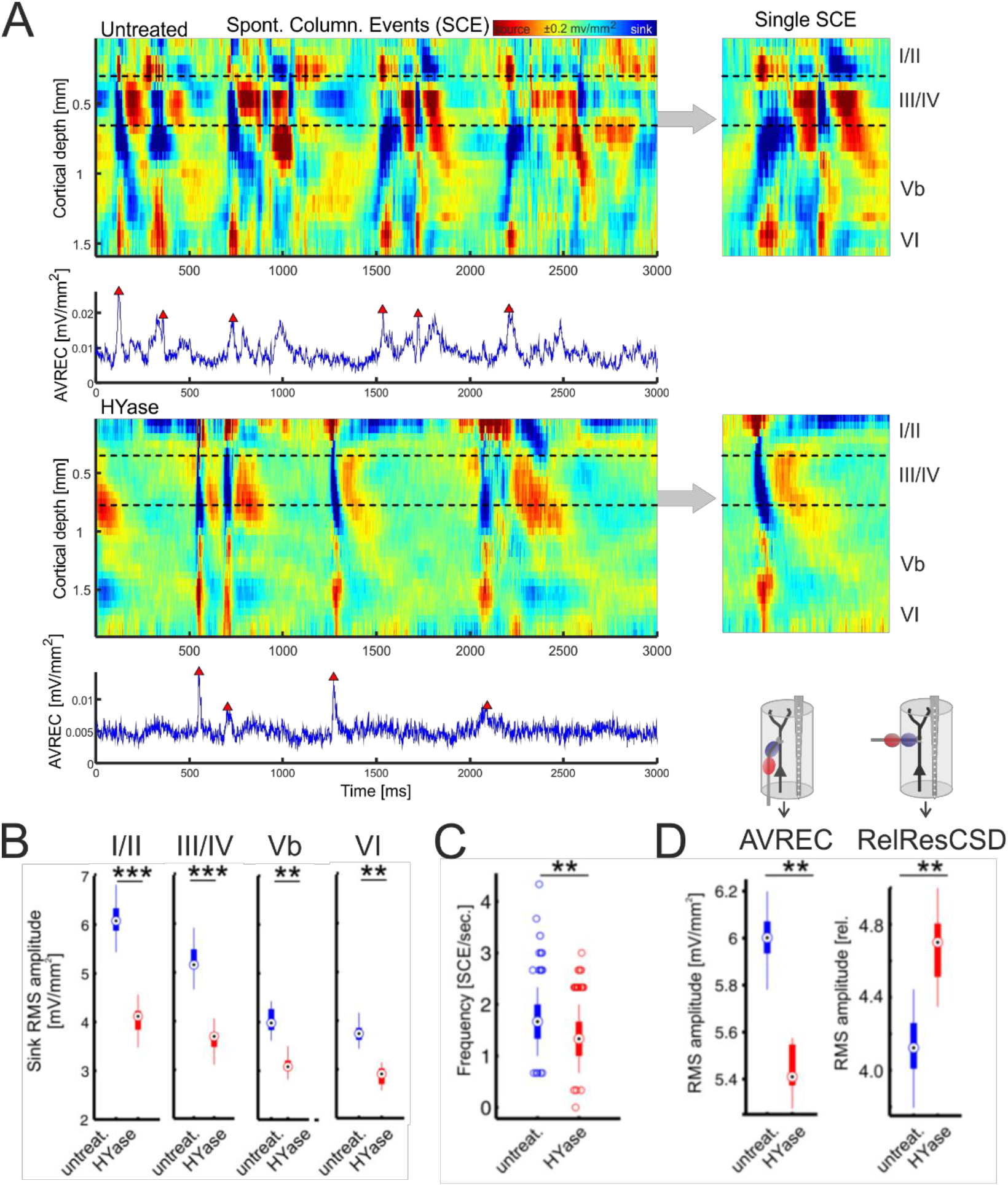
Injection of HYase reduced the frequency of spontaneous columnar events and increased crosscolumnar activity spread. **A.** *Top,* Recordings of spontaneous activity showed the occurrence of spontaneous events of translaminar synaptic information flow mimicking evoked columnar feedforward activation profiles called spontaneous columnar events. SCE’s in untreated cortex showed information flow starting from pacemaker activity in layer Vb (see inlay right). *Bottom*, After Injection of HYase SCE’s were shorter in duration, showed no infragranular pacemaker structure and did not expand through all cortical layers (see inlay right). Frequency of SCE’s was quantified by peak detection of the AVREC trace based on crossings of 3SD above baseline (blue traces). **B.** Mean RMS amplitudes (±SEM) within separate cortical layers were all significantly reduced after HYase treatment. **C.** Frequency of SCE’s per second was significantly reduced after HYase treatment. **D**. Decreased RMS amplitude of the AVREC (±SEM) was paralleled with a significant increase of the RMS amplitude (±SEM) of the RelResCSD. Asterisks mark significant differences between groups (paired Student’s t-test, **p<0.01; ***p<0.001).

## Discussion

This study investigated the impact of acute removal of the extracellular matrix on the network physiology in the gerbil primary auditory cortex by local injection of the ECM-degrading enzyme hyaluronidase. Based on laminar CSD recordings we found that ECM removal altered the spatiotemporal profile of sensory-evoked synaptic population activity across cortical layers. Stronger activation of supragranular layers I/II was paralleled by an increased spectral integration via lateral corticocortical activity spread, while tone-evoked activation in infragranular layer Vb was reduced. That further coincided with increased evoked oscillatory power in the beta band (25-36 Hz) within infragranular layer Vb revealed by a multitaper spectral analysis of layer-specific CSD activity. We then applied time-domain Granger causality measures to map the directional causal relationship between cortical layers. While we found a significant increase in the GC from supragranular layers I/II towards the infragranular layer Vb, the causal drive from deep infragranular layer VI to granular layers III/IV was significantly reduced. We also found a significant increase in the GC from layer Vb towards infragranular layer VI. Thereby, our data shows that enzymatic removal of the ECM in sensory cortex acutely alters the entire columnar synaptic network processing under control of supragranular layer activity. Columnar processing is thereby dominated by the integration of early local synaptic inputs and crosscolumnar widespread activity in upper layers.

### Effects of ECM removal on sensory-evoked activity in auditory cortical circuits

While we generally found increased activity in supragranular layers across all stimulation frequencies, early synaptic inputs in infragranular layers Vb, related to activity of thalamocortical collaterals, were reduced particularly for BF stimulation (Fig. 2A). Early granular layer III/IV input activity was unaffected implying an asymmetric impact of ECM removal on different thalamocortical input systems in sensory cortex (Constantinople and Bruno, 2013). This is in accordance with reports that sensory input in layer Vb is more vulnerable to juvenile forms of plasticity, as induced by monocular deprivation (Medini, 2014). Thalamocortical collaterals terminating in infragranular layer Vb also transmit early subcortical inputs (cf Fig. 1) but might be less responsible for the spectral integration across the frequency range at a given cortical patch. Here, granular circuits in auditory cortex have been related to a local amplification of sensory input via recurrent microcircuits yielding a robust bottom-up driven tonotopic tuning (Liu et al., 2007). The shift of the entire network activity towards upper layers (see also Fig. 2B) might hence represent an altered and layer-specific input function of spectral information after acute ECM removal. We further found that the overall columnar activity, quantified by the AVREC, was increased accompanied by a frequency-specific gain increase in the RelResCSD (Fig. 3). We found a corresponding increase of the response bandwidth of the RelResCSD which indicates increased lateral corticocortical activity spread after ECM removal.

With maturation of cortical circuits and phases of experience-dependent shaping of crosscolumnar connectivity patterns, intracolumnar circuits stabilize subsequently in order to control its outputs to promote behavior later on (Crocker-Buque et al., 2015). In accordance, learning-related plasticity of synaptic circuits during auditory learning has been linked mainly to corticocortical synaptic connections within supragranular cortical layers I/II (Francis et al., 2018; Froemke et al., 2007), while thalamic input layers III/IV and Vb show rather moderate plastic changes. Thus, increase of synaptic integration in supragranular layers in the adult cortex after ECM removal may promote plastic adaptations of corticocortical synaptic circuits during learning (Happel et al., 2014b; Lottem et al., 2018; Niekisch et al., 2019; Romberg et al., 2013). We hypothesize that a relative imbalance of synaptic activity between supragranular layers I/II (mediated by corticocortical inputs) and infragranular layer Vb (mediated by early thalamocortical input) is one of the possible physiological mechanisms by which HYase is mediating its action. To further reveal potential network mechanisms of this newly proposed hypothesis, we continued with an analysis of layer-specific neuronal oscillations and translaminar network dynamics.

### Oscillatory layer-specific dynamics after ECM removal

We found that stimulus-evoked oscillations showed a layer-dependent shift of increased beta power (~25-36 Hz) selectively in infragranular layer Vb (Fig. 4). Infragranular beta oscillations have been linked to altered translaminar processing in line with increased drive from corticocortical inputs in upper layers (Schmiedt et al., 2014; Sherman et al., 2016). In the human EEG and animal LFP literature, beta range fluctuations have been related to the deployment of top-down attention (Bosman et al., 2012; Miller and Buschman, 2013), working memory allocation (Salazar et al., 2012; Tallon-Baudry et al., 2004), and stimulus salience (Maier et al., 2008; Wilke et al., 2006). Increased beta power in deep layers and reverse effects on high-frequency components in upper layers has been specifically linked to altered GABA levels in PV+ interneurons in PV-GAD67 mice (Kuki et al., 2015), suggesting that PV cells, generally rich in ECM, might play an important roles in balancing neuronal oscillations across cortical layers.

Further, Increased layer Vb beta power has further been attributed to relatively increased inputs on distal over proximal dendrites of intrinsic bursting infragranular neurons in upper layers (Sherman et al., 2016). Those neurons therefore might play a pivotal role for orchestrating the translaminar columnar activity. Time-domain Granger causality analysis further supported our hypothesis of upper layers dominating the cortical column activity with an increased causal drive of supragranular layers I/II towards infragranular layer Vb. Further, we found an altered dynamical relationship of deeper cortical layers. While layer VI activity became more strongly dependent on layer Vb input, its output drive onto granular layer III/IV was decreased after ECM removal (Fig. 6). This decrease in feedback drive from infragranular layer VI to granular layer III/IV is important in controlling the thalamocortical input within granular layer III/IV (Lee et al., 2012). Thereby, ECM removal might induce a higher weight of local sensory input processing and its corticocortical integration and reduced activity of effector circuits towards downstream areas, as motor-related cortical outputs (Lee et al., 2008; Voigts et al., 2019; Williamson and Polley, 2019) potentially supporting higher plastic adaptability towards altered sensory inputs.

### ECM removal as a model for cortical “re-juvenation” and adult learning?

ECM removal has been linked to re-open ‘critical periods’ of juvenile plasticity unlocking the potential of considerable topographic map adaptations (Gundelfinger et al., 2010; Pizzorusso et al., 2002). ECM removal therefore has been interpreted as a temporally constricted period of neuronal “re-juvenation” promoting a higher level of structural, developmental plasticity (Berardi et al., 2004; Fawcett et al., 2019b; Lensjø et al., 2017). States of weakened ECM have been also linked to other forms of plasticity, as for instance post-traumatic restoration, recognition memory, seasonal learning in songbirds, and cognitive flexibility during reversal learning (Banerjee et al., 2017; Cornez et al., 2020; Happel et al., 2014b; Romberg et al., 2013). Further, mouse models deficient in specific matrix components, as tenascin-R or brevican, have shown impairments in hippocampus-dependent spatial or contextual acquisition learning (Dityatev et al., 2010). Recently, we showed in wildtype mice that brevican was downregulated 48h after auditory training in the auditory cortex (Niekisch et al., 2019). Recently, we further showed that such intrinsic reduction of the ECM integrity in the adult cortex can be linked to the stimulation of D1 dopamine receptors suggesting an activity-dependent and learning-related cellular mechanism (Mitlöhner et al., 2020). Altogether, those studies enunciate the major impact of the integrity of the cortical ECM onto the delicate balance of plasticity and tenacity in the learning brain. However, the underlying physiological circuit mechanisms are still rather elusive.

In this study, we investigated the effects of ECM removal acutely and under ketamine-xylazine anaestesia. We have recently shown that, in addition to generally very similar spatiotemporal activity patterns, anesthesia mainly influences gain enhancement in layer III/IV at best frequency stimulation (Deane et al., 2020). Particularly, ketamine-xylazine had only minor effects on the spectral beta range and off-BF-evoked activity, which showed strongest effects after HYase treatment. In this study, ECM removal specifically enhanced beta power in infragranular layer Vb, and minor effects on layer III/IV activity.

Beta power is also higher in children than adults (Vanvooren et al., 2015), which we hence found to be reversed after “re-juvenation” by HYase (Fig. 5). Furthermore, the reduction in spontaneous columnar events during states of weakened ECM might mimic decreased states of spontaneous synchronicity in juvenile cortex (Luhmann et al., 2016), where peripheral stimulation dominates temporal discharge patterns (Yang et al., 2009). Spontaneous columnar activity in Acx in anaesthetized animals has been linked to a dominant drive of pyramidal neurons and interneurons in deeper layers (Sakata and Harris, 2009). We suppose that ECM removal altered the balance of activity between these cell types reflected in fewer spontaneous translaminar events, but higher crosscolumnar spread compared to the untreated adult cortex. The function of earliest activity in sensory cortex is to establish topographic maps based on experience-dependent plasticity of sensory bottom-up input via thalamocortical circuits (Chiu and Weliky, 2001; Kilb et al., 2011). Early sensory-derived synaptic plasticity is hence pronounced in cortical layers II – IV, but less affecting infragranular synapses (Medini, 2011).

It has been argued before that long-range corticocortical processing in upper layers of the juvenile visual cortex is necessary to correlate activity across cortical space in order to promote such early bottom-up-driven sensory-based plasticity (Chiu and Weliky, 2001; Ko et al., 2013; Luhmann et al., 1990). Consistently, two consecutive days of strabismus cause selective strengthening of horizontal connections in layers II/III between cortical columns representing the same eye (Trachtenberg and Stryker, 2001). The acute cortical state after ECM weakening in our study may resemble such a processing mode by the increase of relative residual contributions of the spontaneous CSD patterns indicating increase of corticocortical activity spread. Further research in awake, behaving animals would be needed to link such circuit activity after removal of the ECM in sensory cortex directly to the unlocking of plastic adaptations of layer-specific network dynamics in the adult brain (cf. Fig. 6). Most importantly, behavioral effects of ECM modulations are most prominent after a certain time of training (Happel et al., 2014b; Romberg et al., 2013), which let us suppose, that they may have no immediate impact on perception and behavior. Nevertheless, we have unraveled a potential circuit mechanism, that potentially allows the cortical network to adjust the balance of internal activity and sensory-driven inputs (Toyoizumi et al., 2013). Ultimately, such changes may promote developmental (Gundelfinger et al., 2010) and learning-derived plasticity underlying increased performance in cognitively demanding tasks and memory recall over time (Fawcett et al., 2019b; Frischknecht et al., 2017; Patton et al., 2019).

## Methods and Materials

### Experimental Design

Experiments were performed on 12 adult male ketamine-xylazine anesthetized Mongolian gerbils (age: 4-10 months, body weight: 80-100 g; n=9 HYase injection, n=3 control injections). Surgical and experimental procedures have been described in detail previously (Brunk et al., 2019; Happel et al., 2010; 2014). All experiments were conducted in accordance with the international NIH Guidelines for Animals in Research and with ethical standards for the care and use of animals in research defined by the German Law for the protection of experimental animals. Experiments were approved by an ethics committee of the state Saxony-Anhalt, Germany.

### Surgery and electrophysiological recordings

Mongolian gerbils (n=12) were anesthetized by intraperitoneal infusion (0.06ml/h) of 45% ketamine (50 mg/ml, Ratiopharm, Germany), 5% xylazine (Rompun, 2%, BayerVital, Germany) and 50% isotonic sodium chloride solution (154mmol/l, Braun, Germany). Status of anesthesia was monitored, and body temperature was kept at 37°C. Right ACx was exposed by craniotomy (~3×4mm) of the temporal bone. Recordings were performed in an acoustically and electrically shielded recording chamber. Laminar profiles of local field potentials (LFP) were measured using linear 32-channel-shaft electrodes (NeuroNexus, 50 μm inter-channel spacing, 413 μm2 site area; type A1×32-5mm-50-413) inserted perpendicular to the cortical surface. Neuronal potentials were preamplified (500x), band-pass filtered between 0.7-170 Hz (3 dB cut-off frequency), digitized at 2 kHz (Multichannel Acquisition Processor, Plexon Inc.) and averaged over 40-80 stimulus repetitions. The location of the field AI in primary ACx was identified by vasculature landmarks and physiological parameters (Happel et al., 2010). In this study, we analyzed only data from recordings which exceed 90 minutes after implantation of the electrode–a state where the cortical responses under anesthesia have been largely stabilized as reported recently (Deane et al., 2020).

### Enzymatic removal of the extracellular matrix by intracortical injections

In direct proximity of the recording electrode, we inserted a 3.5-inches glass capillary (<250 μm distance) approximately within the middle of the ACx in depths of 500 μm in order to perform microinjections (Nanoliter injector 2000, WPI) of HYase at the cortical patch of recording in ACx. We first recorded baseline activity in order to reveal stabilized cortical responses after insertion of micropipette and electrode. Then, we recorded spontaneous and tone-evoked activity over a period of roughly 1h. After that, we injected 500 nl of HYase-solution (500 units) in 22 steps (22.8 nl each) with 3 seconds pause in between (Happel et al., 2014b). Before we continued recordings, we waited at least 2.5h until we continued with recordings after ECM removal.

### Current source density and residue analysis

One-dimensional current-source density (CSD) profiles were calculated from the second spatial derivative of the LFP (Mitzdorf, 1985; Schroeder et al., 1998):

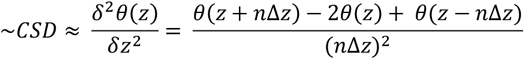

where *θ* is the field potential, z the spatial coordinate perpendicular to the cortical laminae, *Δz* the spatial sampling interval (50 μm), and *n* the differentiation grid. LFP profiles were smoothed with a weighted average (Hamming window) of 7 channels (corresponding to a spatial filter kernel of 300 μm; linear extrapolation of 4 channels at boundaries; see Happel et al., 2010). Main sink components were found to represent the architecture of primary sensory input from the lemniscal auditory part of the thalamus –the medial geniculate body (MGB). Lemniscal thalamocortical projections from the ventral division of the MGB terminate on pyramidal neurons with local dendritic arbors and ramifying axons in cortical layers III/IV, which are henceforth referred to as the granular inputs layers. Collaterals of these thalamocortical projections also target infragranular layer Vb (Constantinople and Bruno, 2013; Schaefer et al., 2015; Winer et al., 2005). Layers above (I/II) or beyond (V-VI) the granular layers are generally referred to as supragranular and infragranular layers, respectively (Winer, 1984). In our study, corresponding sink activity within cortical layers were referred to as granular activity (III/IV) supragranular activity (layer I/II), and infragranular activity (layer Vb and VI). Rout-mean square (RMS) amplitudes of individual layers were determined for individual channels and then averaged.

Based on single trial CSD profiles without spatial filtering we transformed the CSD by rectifying and averaging waveforms of each channel (n) comprising the laminar CSD profile (AVREC). The AVREC waveform provides a measure of the temporal pattern of the overall strength of transmembrane current flow (Schroeder et al., 1998). The relative residue of the CSD (RelResCSD), defined as the sum of the non-rectified divided by the rectified magnitudes for each channel, was used to quantify the balance of the transmembrane charge transfer along the recording axis (Happel et al., 2010):

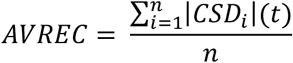

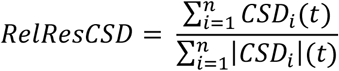

We have shown before that the RelResCSD provides a quantitative measure of the contributions of intercolumnar, corticocortical synaptic circuits to the overall measured local cortical activity. In short, the rationale is that corticocortical circuits would be more likely distributed beyond the integration cylinder surrounding the electrode array in which extracellular currents contribute to the measured LFP (cf. Fig. 3). We quantified the RMS values of both parameters during the period of stimulus presentation (200 ms).

### Auditory stimulation and estimation of tuning curves

We further presented pseudo-randomized series of pure tones (200 ms with 5 ms sinusoidal rising and falling ramps; spanning 8 octaves from 250 Hz to 32 kHz; 40 dB over threshold; inter-stimulus interval: 600 ms; digitally synthesized using Matlab and converted to analog signals by a data acquisition National Instruments card; PCI-BNC2110). Stimuli were delivered via a programmable attenuator (g.PAH, Guger Technologies; Austria), a wide-range audio amplifier (Thomas Tech Amp75) and a loudspeaker (Tannoy arena satellite) positioned in 1m distance to the animal’s head. The response threshold was determined as the lowest intensity eliciting a significant response at any frequency 2SD over baseline (> 5ms). The sound level then was adjusted to 40dB above threshold, which generally was at intensities around 65 dB SPL. Response bandwidths of AVREC and RelResCSD were quantified as the Q40dB-values corresponding to the number of octaves evoking a tone-evoked response over the threshold criterion (Happel and Ohl, 2017). The best frequency (BF) of the recording site was determined as the frequency evoking the highest RMS amplitude within the granular layer III/IV at 40dB above threshold. For estimation of pure-tone based frequency response curves, evoked RMS amplitudes of individual parameters were binned into frequency bins with respect to their distance to the best frequency of each individual recording (─3-4 oct.; ─1-2 oct.; BF; +1-2 oct.; +3-4 oct.). Cross-laminar activation patterns were quantified and statistically analyzed by introducing a Layer-Symmetry-Index (LSI). The LSI quantifies the total activity bias based on RMS values of the supragranular layer vs. infragranular layer Vb by calculating LSI = (I/II – Vb) / (I/II + Vb). A positive LSI indicates that columnar synaptic activity is biased towards stronger supragranular activity, while negative values indicate stronger recruitment of infragranular synaptic circuits.

### Recording and analysis of spontaneous activity

We recorded spontaneous data in 7 animals for at least 3 minutes before and after injection of HYase. Spontaneous CSD data was cut into 6 s traces and we analyzed RMS values of the AVREC, RelResCSD and layer dependent CSD traces of individual sink components (see above). Spontaneous events of translaminar synaptic information were referred to as spontaneous columnar events (SCE). The frequency of SCE’s was quantified by a peak detection of the AVREC trace based on crossings of 3SD above baseline (median value of the whole spontaneous trace) and at least 150ms between individual peaks.

### Spectral analysis of CSD data

Traditional Fourier spectral analysis uses a single windowing function, which yields large variance and biases especially at higher frequencies. We therefore used the multitaper Fourier transform employing multiple splenial tapers that are orthogonal to each other (Chronux toolbox; http://chronux.org), as explained in detail elsewhere (Deliano et al., 2018). CSD-signals were transformed into the frequency domain by a multitaper FFT (600 ms epochs) and 5 tapers of 600 ms length with a time-bandwidth product of 3 and no padding were employed (Bokil et al., 2010). The stimulus evoked power spectrum was averaged by the complex valued spectra across trials before calculating the power as squared magnitude of the average. To normalize the power spectral values, we divided it by the sum of all values. As induced spectra might include background effects unrelated to the stimulus, we further calculated the power spectrum during spontaneous recordings. Finally, the mean and the standard error of the mean of the power spectra were calculated across subjects. Power spectral comparison before and after HYase administration was done across subjects, separately for the BF condition and layer (I/II, III/IV, Vb, VI). Comparisons were restricted to a frequency range from 1 to 100 Hz. To avoid biased determination of power spectrum bands, t-scores were calculated between recordings in untreated cortex and after HYase injection for each spectrum, frequency, and layer using sample mean and variance and the significance level was corrected using a Benjamin Hochberg procedure as a suitable method to adjust the false discovery rate when performing a large number of multiple comparisons (Benjamini and Hochberg, 2000). Accordingly, we ranked individual p-values in an ascending order. Critical Benjamin-Hochberg (BH) p-values are calculated by p*= (i_r_ / m)*Q, where i_r_ is the individual p-value’s rank, m is the total number of tests, and Q is the false discovery rate (0.1). The critical BH p* value serves as adapted level of significance.

### Time-domain conditional granger causality

To identify the dynamical nature of the directed connectivity between different cortical layers as defined by the CSD analysis, we calculated the multivariate conditional causality in the time-domain using MVGC Matlab toolbox (Barnett and Seth, 2014). Generally, unconditional Granger causality (GC) assumes two time series (X, Y) and provides information about the magnitude of their causal dependency. Briefly, the unconditional GC measures if past values of the time series Y provide information that predicts upcoming values X better than the past history of X itself. Mathematically, this is done by constructing and comparing two linear vector autoregressive models (VAR). the first model described as a full model where the value of X at particular time point is a linear weighted sum of both its own and Y’s past values according to the following equation:

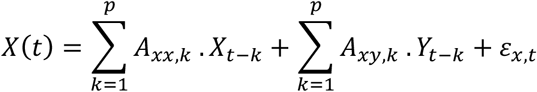

where p is the model order, *A*_*xx*,*k*_ and *A*_*xy*,*k*_ are the full regression coefficient matrices and *ε*_*x,t*_ is the error or residual process. The second model is described as the reduced model where we only use X past values to predict X current values.

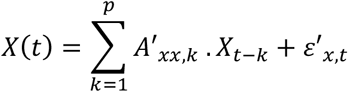

By taking the logarithmic ratio between the covariance of the two residuals we can get the estimate F describing Y being G-causal of X.

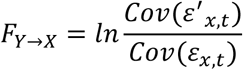

In other words, G-causal quantifies the reduction in the prediction error when including Y past values in the model. The main limitation of the unconditional GC is the spurious causal relationship that can be derived if exist a third variable Z that both X and Y depends on. the conditional GC takes into account the inter-dependent nature of the jointly distributed multivariate times series data by including the variables Z in the reduced and full regression model and now calculating the conditional G-causal F between X, Y is explained as the following

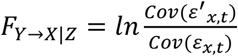

The multivariate granger causality toolbox adds to the standard G-causal estimation as it allows for reduced model parameter computation from the full model parameters for more computationally effective and accurate statistical estimates (Barnett and Seth, 2014). We used the toolbox to estimate the conditional granger causality from the single trial evoked CSD dataset. To ensure that our data was stationary, we took the first time-derivative of CSD traces and then tested for stationarity using the Augmented Dickey Fuller test. After that, we estimated the order of our vector autoregressive (VAR) model using Akiko information criteria (AIC). The model order is then used to estimate the model parameters (the regression coefficients and residual errors) using the “LWR algorithm” or the multivariate extensions to Durbin recursion. The autocovariance sequence derived from the full model parameters is estimated and used to derive the reduced VAR parameters using an algorithm to solve the Yule-Walker equation. The residual covariance matrices of both the reduced and full model is used to estimate the G-causal as in equation 7. the G-causal estimates for each animal is then log transformed, and a ratio paired t-test is performed. A false discovery rate of 0.05 was set for multiple comparison and a critical p-value is determined as a cut off for significance (Benjamini and Hochberg, 2000).

### Immunohistochemistry

To assess effects of HYase injection on the ECM, we applied microinjections in naïve mature gerbils (≥3 month; n=4) under anesthesia as explained above (5 mg ketamine and 3 mg xylazine per 100 g body weight, i.p.). We injected HYase in the ACx unilaterally and used 0.9% saline at the opposite side as a control. Animals were transcardially perfused with 20 ml phosphate buffered saline (PBS) followed by 200 ml paraformaldehyde (4% in PBS) 2.5h after injections in order to assess the ECM removal at the earliest time point where recordings have been done in the experimental animals. Brains were removed, postfixated overnight, cryoprotected in 30% sucrose solution for 48h, frozen, and cut in a series of frontal sections (40μm thickness) on a cryostat (Leica CM 1950; Germany). After blocking (5% goat serum) and washing, the sections were stained using *Wisteria floribunda* agglutinin coupled to fluorescein (WFA; 1:100; cat# FL-1351, RRID: AB_2336875, Vector Laboratories, CA, USA) for 12h (at 4°C). Then, sections were incubated with anti-PV (mouse, 1:4000, 12h, cat# 235, RRID: AB_10000343, Swant, Switzland) and afterwards with a Cy3-coupled anti-mouse secondary antibody (goat, 1:500, 2h, cat#115-165-166, RRID: AB_2491007, Dianova, Germany). Finally, sections were mounted on gelatin-coated slides and coverslipped with MOWIOL (Fluka, Germany). Images were taken using a confocal microscope (Zeiss LSM 700, Germany) equipped with a 2.5x objective (NA 0.085, Zeiss, Germany).

### Statistical analysis

Comparison of multiple groups was performed by multifactorial repeated-measures ANOVAs (rmANOVAs). For comparison between two groups, paired sample Student’s t tests were used. Generally, a significance level of α = 0.05 was chosen. For multiple comparisons, we Bonferroni-corrected levels of significance.

**The authors declare no conflict of interest**

## Acknowledgements

The work was supported by grants from the Deutsche Forschungsgemeinschaft DFG (SFB779) and the Leibniz-Society WGL granted to RF and MFKH.

## Author contributions

MT and MFKH conceived the study and designed the experiments. Experiments and data analysis was carried out by MT and supervised by MFKH. MD developed statistical test procedures. JH and EB performed histological experiments. RF supplied agents. Figures were prepared by MT and MFKH. MT and MFKH wrote the paper with the assistance from all co-authors.

## Supplemental Information

**Supplemental Fig. 1.**
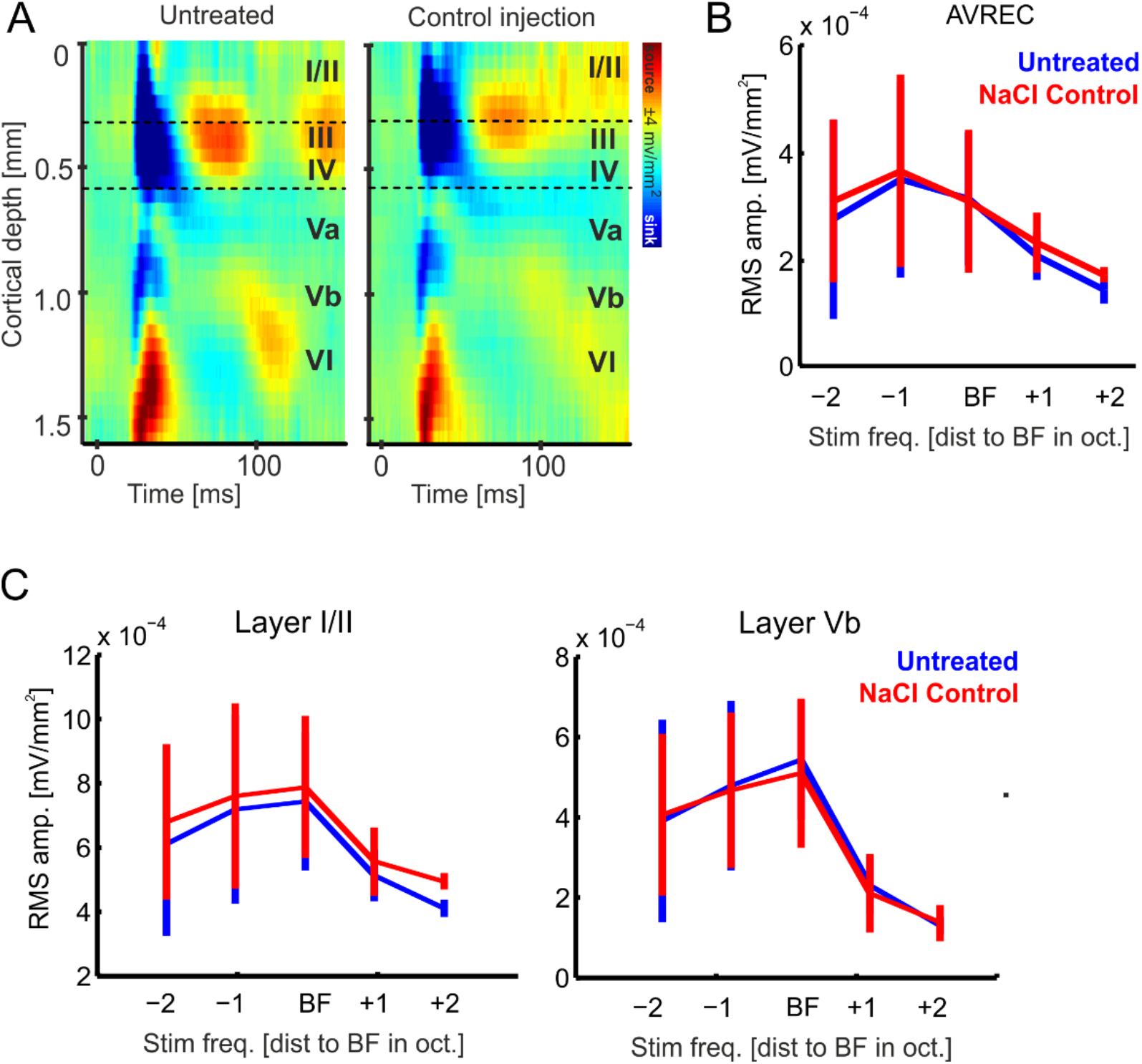
Control injection of 0.9% sodium-chloride (NaCl) did not change layer-specific processing in the auditory cortex. **A.** Tone-evoked CSD profiles before and after control injection of 0.9% sodium-chloride displayed the canonical feedforward pattern of early and late sink activity distributed across cortical layers. Control injections did not lead to qualitative changes of tone-evoked activity. **B**. Frequency-response tuning curves of AVREC RMS amplitudes (±SEM) showed no significant change of activity across all stimulation frequencies after control NaCl injection **C.** Frequency-response tuning curves for the RMS amplitude (±SEM) of CSD traces from cortical layers I/II and Vb (which were altered after ECM removal) did also not show any significant change after control injections. Statistical significance was tested by a 2-way rmANOVA with factors ‘Freq’ and ‘Injection’. All main effects and interactions were not significant.

## Literature

Banerjee SB, Gutzeit VA, Baman J, Aoued HS, Doshi NK, Liu RC, Ressler KJ. 2017. Perineuronal Nets in the Adult Sensory Cortex Are Necessary for Fear Learning. Neuron 95:169–179.e3. doi:10.1016/j.neuron.2017.06.007

Barnett L, Seth AK. 2014. The MVGC multivariate Granger causality toolbox: A new approach to Granger-causal inference. J Neurosci Methods. doi:10.1016/j.jneumeth.2013.10.018

Benjamini Y, Hochberg Y. 2000. On the adaptive control of the false discovery rate in multiple testing with independent statistics. J Educ Behav Stat. doi:10.3102/10769986025001060

Berardi N, Pizzorusso T, Maffei L. 2004. Extracellular matrix and visual cortical plasticity: Freeing the synapse. Neuron 44:905–908. doi:10.1016/j.neuron.2004.12.008

Bokil H, Andrews P, Kulkarni JE, Mehta S, Mitra PP. 2010. Chronux: A platform for analyzing neural signals. J Neurosci Methods. doi:10.1016/j.jneumeth.2010.06.020

Bosman C a., Schoffelen JM, Brunet N, Oostenveld R, Bastos AM, Womelsdorf T, Rubehn B, Stieglitz T, De Weerd P, Fries P. 2012. Attentional Stimulus Selection through Selective Synchronization between Monkey Visual Areas. Neuron 75:875–888. doi:10.1016/j.neuron.2012.06.037

Brunk MGK, Deane KE, Kisse M, Deliano M, Vieweg S, Ohl FW, Lippert MT, Happel MFK. 2019. Optogenetic stimulation of the VTA modulates a frequency-specific gain of thalamocortical inputs in infragranular layers of the auditory cortex. Sci Rep 9:https://doi.org/10.1038/s41598-019-56926-6. doi:10.1101/669168

Chiu C, Weliky M. 2001. Spontaneous activity in developing ferret visual cortex in vivo. J Neurosci 21:8906–14.

Constantinople CM, Bruno RM. 2013. Deep cortical layers are activated directly by thalamus. Science 340:1591–4. doi:10.1126/science.1236425

Cornez G, Collignon C, Müller W, Ball GF, Cornil CA, Balthazart J. 2020. Seasonal changes of perineuronal nets and song learning in adult canaries (Serinus canaria). Behav Brain Res. doi:10.1016/j.bbr.2019.112437

Crocker-Buque A, Brown SM, Kind PC, Isaac JTRR, Daw MI. 2015. Experience-Dependent, Layer-Specific Development of Divergent Thalamocortical Connectivity. Cereb Cortex 25:2255–2266. doi:10.1093/cercor/bhu031

de Vivo L, Landi S, Panniello M, Baroncelli L, Chierzi S, Mariotti L, Spolidoro M, Pizzorusso T, Maffei L, Ratto GM. 2013. Extracellular matrix inhibits structural and functional plasticity of dendritic spines in the adult visual cortex. Nat Commun 4:1484. doi:10.1038/ncomms2491

Deane KE, Brunk MGK, Curran AW, Zempeltzi MM, Ma J, Lin X, Abela F, Aksit S, Deliano M, Ohl FW, Happel MFK. 2020. Ketamine anesthesia induces gain enhancement via recurrent excitation in granular input layers of the auditory cortex. J Physiol. doi:10.1113/jp279705

Deliano M, Brunk MGK, El-Tabbal M, Zempeltzi MM, Happel MFK, Ohl FW. 2018. Dopaminergic neuromodulation of high gamma stimulus phase-locking in gerbil primary auditory cortex mediated by D1/D5-receptors. Eur J Neurosci. doi:10.1111/ejn.13898

Dick G, Liktan C, Alves JN, Ehlert EME, Miller GM, Hsieh-Wilson LC, Sugahara K, Oosterhof A, Van Kuppevelt TH, Verhaagen J, Fawcett JW, Kwok JCF. 2013. Semaphorin 3A binds to the perineuronal nets via chondroitin sulfate type E motifs in rodent brains. J Biol Chem 288:27384–27395. doi:10.1074/jbc.M111.310029

Dityatev A, Schachner M, Sonderegger P. 2010. The dual role of the extracellular matrix in synaptic plasticity and homeostasis. Nat Rev Neurosci 11:735–746. doi:10.1038/nrn2898

Fawcett JW, Oohashi T, Pizzorusso T. 2019a. The roles of perineuronal nets and the perinodal extracellular matrix in neuronal function. Nat Rev Neurosci 20:451–465. doi:10.1038/s41583-019-0196-3

Fawcett JW, Oohashi T, Pizzorusso T. 2019b. The roles of perineuronal nets and the perinodal extracellular matrix in neuronal function. Nat Rev Neurosci 20:451–465. doi:10.1038/s41583-019-0196-3

Francis NA, Elgueda D, Englitz B, Fritz JB, Shamma SA. 2018. Laminar profile of task-related plasticity in ferret primary auditory cortex. Sci Rep. doi:10.1038/s41598-018-34739-3

Frischknecht R, Happel MFK, Happel MFK. 2017. Impact of the extracellular matrix on plasticity in juvenile and adult brains. e-Neuroforum 22:1–6. doi:10.1515/s13295-015-0021-z

Frischknecht R, Heine M, Perrais D, Seidenbecher CI, Choquet D, Gundelfinger ED. 2009. Brain extracellular matrix affects AMPA receptor lateral mobility and short-term synaptic plasticity. Nat Neurosci 12:897–904. doi:10.1038/nn.2338

Frischknecht R, Seidenbecher CI. 2012. Brevican: A key proteoglycan in the perisynaptic extracellular matrix of the brain. Int J Biochem Cell Biol 44:1051–1054. doi:10.1016/j.biocel.2012.03.022

Froemke RC, Merzenich MM, Schreiner CE. 2007. A synaptic memory trace for cortical receptive field plasticity. Nature 450:425–429. doi:10.1038/nature06289

Gogolla N, Caroni P, Lüthi A, Herry C. 2009. Perineuronal nets protect fear memories from erasure. Science (80-) 325:1258–1261. doi:10.1126/science.1174146

Gundelfinger ED, Frischknecht R, Choquet D, Heine M. 2010. Converting juvenile into adult plasticity: A role for the brain’s extracellular matrix. Eur J Neurosci 31:2156–2165. doi:10.1111/j.1460-9568.2010.07253.x

Happel MFK, Deliano M, Handschuh J, Ohl FW. 2014a. Dopamine-modulated recurrent corticoefferent feedback in primary sensory cortex promotes detection of behaviorally relevant stimuli. J Neurosci 34:1234–47. doi:10.1523/JNEUROSCI.1990-13.2014

Happel MFK, Jeschke M, Ohl FW. 2010. Spectral integration in primary auditory cortex attributable to temporally precise convergence of thalamocortical and intracortical input. J Neurosci 30:11114–27. doi:10.1523/JNEUROSCI.0689-10.2010

Happel MFK, Niekisch H, Castiblanco Rivera LL, Ohl FW, Deliano M, Frischknecht R. 2014b. Enhanced cognitive flexibility in reversal learning induced by removal of the extracellular matrix in auditory cortex. Proc Natl Acad Sci U S A 111:2800–2805. doi:10.1073/pnas.1310272111

Happel MFK, Ohl FW. 2017. Compensating Level-Dependent Frequency Representation in Auditory Cortex by Synaptic Integration of Corticocortical Input. PLoS One 12:e0169461. doi:10.1371/journal.pone.0169461

Heine M, Groc L, Frischknecht R, Béïque J-CC, Lounis B, Rumbaugh G, Huganir RL, Cognet L, Choquet D. 2008. Surface mobility of postsynaptic AMPARs tunes synaptic transmission. Science (80-) 320:201–5. doi:10.1126/science.1152089

Hylin MJ, Orsi S a, Moore AN, Dash PK. 2013. Disruption of the perineuronal net in the hippocampus or medial prefrontal cortex impairs fear conditioning. Learn Mem 20:267–73. doi:10.1101/lm.030197.112

Kaur S, Lazar R, Metherate R. 2004. Intracortical pathways determine breadth of subthreshold frequency receptive fields in primary auditory cortex. J Neurophysiol 91:2551–67. doi:10.1152/jn.01121.2003

Kilb W, Kirischuk S, Luhmann HJ. 2011. Electrical activity patterns and the functional maturation of the neocortex. Eur J Neurosci 34:1677–1686. doi:10.1111/j.1460-9568.2011.07878.x

Ko H, Cossell L, Baragli C, Antolik J, Clopath C, Hofer SB, Mrsic-Flogel TD. 2013. The emergence of functional microcircuits in visual cortex. Nature 496:96–100. doi:10.1038/nature12015

Kuki T, Fujihara K, Miwa H, Tamamaki N, Yanagawa Y, Mushiake H. 2015. Contribution of parvalbumin and somatostatin-expressing GABAergic neurons to slow oscillations and the balance in beta-gamma oscillations across cortical layers. Front Neural Circuits 9:6. doi:10.3389/fncir.2015.00006

Lee CC, Lam YW, Sherman SM. 2012. Intracortical convergence of layer 6 neurons. Neuroreport. doi:10.1097/WNR.0b013e328356c1aa

Lee SH, Carvell GE, Simons DJ. 2008. Motor modulation of afferent somatosensory circuits. Nat Neurosci. doi:10.1038/nn.2227

Lensjø KK, Lepperød ME, Dick G, Hafting T, Fyhn M. 2017. Removal of perineuronal nets unlocks juvenile plasticity through network mechanisms of decreased inhibition and increased gamma activity. J Neurosci 37:1269–1283. doi:10.1523/JNEUROSCI.2504-16.2016

Liu B, Wu GK, Arbuckle R, Tao HW, Zhang LI. 2007. Defining cortical frequency tuning with recurrent excitatory circuitry. Nat Neurosci 10:1594–600. doi:10.1038/nn2012

Lottem E, Banerjee D, Vertechi P, Sarra D, Lohuis MO, Mainen ZF. 2018. Activation of serotonin neurons promotes active persistence in a probabilistic foraging task. Nat Commun 9:1–12. doi:10.1038/s41467-018-03438-y

Luhmann HJ, Greuel JM, Singer W. 1990. Horizontal Interactions in Cat Striate Cortex: II. A Current Source-Density Analysis. Eur J Neurosci 2:358–368. doi:10.1111/j.1460-9568.1990.tb00427.x

Luhmann HJ, Sinning A, Yang J, Reyes-puerta V. 2016. Spontaneous Neuronal Activity in Developing Neocortical Networks : From Single Cells to Large-Scale Interactions. Front Neural Circuits 10:1–14. doi:10.3389/fncir.2016.00040

Maier A, Wilke M, Aura C, Zhu C, Ye FQ, Leopold DA. 2008. Divergence of fMRI and neural signals in V1 during perceptual suppression in the awake monkey. Nat Neurosci. doi:10.1038/nn.2173

Medini P. 2014. Experience-dependent plasticity of visual cortical microcircuits. Neuroscience 278:367–384. doi:10.1016/j.neuroscience.2014.08.022

Medini P. 2011. Layer- and cell-type-specific subthreshold and suprathreshold effects of long-term monocular deprivation in rat visual cortex. J Neurosci 31:17134–48. doi:10.1523/JNEUROSCI.2951-11.2011

Miller EK, Buschman TJ. 2013. Cortical circuits for the control of attention. Curr Opin Neurobiol. doi:10.1016/j.conb.2012.11.011

Mitlöhner J, Kaushik R, Niekisch H, Blondiaux A, Gee CE, Happel MFK, Gundelfinger E, Dityatev A, Frischknecht R, Seidenbecher C. 2020. Dopamine Receptor Activation Modulates the Integrity of the Perisynaptic Extracellular Matrix at Excitatory Synapses. Cells 9:260. doi:10.3390/cells9020260

Mitzdorf U. 1985. Current source-density method and application in cat cerebral cortex: investigation of evoked potentials and EEG phenomena. Physiol Rev 65:37–100.

Niekisch H, Steinhardt J, Berghäuser J, Bertazzoni S, Kaschinski E, Kasper J, Kisse M, Mitlöhner J, Singh JB, Weber J, Frischknecht R, Happel MFK. 2019. Learning Induces Transient Upregulation of Brevican in the Auditory Cortex during Consolidation of Long-Term Memories. J Neurosci 39:7049–7060. doi:10.1523/jneurosci.2499-18.2019

Pandya NJ, Seeger C, Babai N, Gonzalez-Lozano MA, Mack V, Lodder JC, Gouwenberg Y, Mansvelder HD, Danielson UH, Li KW, Heine M, Spijker S, Frischknecht R, Smit AB. 2018. Noelin1 Affects Lateral Mobility of Synaptic AMPA Receptors. Cell Rep. doi:10.1016/j.celrep.2018.06.102

Patton MH, Blundon JA, Zakharenko SS. 2019. Rejuvenation of plasticity in the brain: opening the critical period. Curr Opin Neurobiol 54:83–89. doi:10.1016/j.conb.2018.09.003

Pizzorusso T, Medini P, Berardi N, Chierzi S, Fawcett JW, Maffei L. 2002. Reactivation of ocular dominance plasticity in the adult visual cortex. Science 298:1248–51. doi:10.1126/science.1072699

Romberg C, Yang S, Melani R, Andrews MR, Horner AE, Spillantini MG, Bussey TJ, Fawcett JW, Pizzorusso T, Saksida LM. 2013. Depletion of perineuronal nets enhances recognition memory and long-term depression in the perirhinal cortex. J Neurosci 33:7057–65. doi:10.1523/JNEUROSCI.6267-11.2013

Sakata S, Harris KD. 2009. Laminar Structure of Spontaneous and Sensory-Evoked Population Activity in Auditory Cortex. Neuron 64:404–418. doi:10.1016/j.neuron.2009.09.020

Salazar RF, Dotson NM, Bressler SL, Gray CM. 2012. Content-specific fronto-parietal synchronization during visual working memory. Science (80-). doi:10.1126/science.1224000

Schaefer MK, Hechavarría JC, Kössl M. 2015. Quantification of mid and late evoked sinks in laminar current source density profiles of columns in the primary auditory cortex. Front Neural Circuits. doi:10.3389/fncir.2015.00052

Schmiedt JT, Maier A, Fries P, Saunders RC, Leopold DA, Schmid MC. 2014. Beta Oscillation Dynamics in Extrastriate Cortex after Removal of Primary Visual Cortex. J Neurosci 34:11857–11864. doi:10.1523/JNEUROSCI.0509-14.2014

Schroeder CE, Mehta a D, Givre SJ. 1998. A spatiotemporal profile of visual system activation revealed by current source density analysis in the awake macaque. Cereb Cortex 8:575–92. doi:10.1093/cercor/8.7.575

Schweitzer B, Singh J, Fejtova A, Groc L, Heine M, Frischknecht R. 2017. Hyaluronic acid based extracellular matrix regulates surface expression of GluN2B containing NMDA receptors. Sci Rep 7. doi:10.1038/s41598-017-07003-3

Sherman MA, Lee S, Law R, Haegens S, Thorn CA, Hämäläinen MS, Moore CI, Jones SR. 2016. Neural mechanisms of transient neocortical beta rhythms: Converging evidence from humans, computational modeling, monkeys, and mice. Proc Natl Acad Sci 113:E4885–E4894. doi:10.1073/pnas.1604135113

Tallon-Baudry C, Mandon S, Freiwald WA, Kreiter AK. 2004. Oscillatory synchrony in the monkey temporal lobe correlates with performance in a visual short-term memory task. Cereb Cortex. doi:10.1093/cercor/bhh031

Toyoizumi T, Miyamoto H, Yazaki-Sugiyama Y, Atapour N, Hensch TK, Miller KD. 2013. A Theory of the Transition to Critical Period Plasticity: Inhibition Selectively Suppresses Spontaneous Activity. Neuron 80:51–63. doi:10.1016/j.neuron.2013.07.022

Trachtenberg JT, Stryker MP. 2001. Rapid anatomical plasticity of horizontal connections in the developing visual cortex. J Neurosci. doi:10.1523/jneurosci.21-10-03476.2001

Vanvooren S, Hofmann M, Poelmans H, Ghesquière P, Wouters J. 2015. Theta, beta and gamma rate modulations in the developing auditory system. Hear Res 327:153–162. doi:10.1016/j.heares.2015.06.011

Voigts J, Deister CA, Moore CI. 2019. Layer 6 ensembles can selectively regulate the behavioral impact and layer-specific representation of sensory deviants. bioRxiv 657114. doi:10.1101/657114

Wilke M, Logothetis NK, Leopold DA. 2006. Local field potential reflects perceptual suppression in monkey visual cortex. Proc Natl Acad Sci U S A. doi:10.1073/pnas.0604673103

Williamson RS, Polley DB. 2019. Parallel pathways for sound processing and functional connectivity among layer 5 and 6 auditory corticofugal neurons. Elife 8:1–21. doi:10.7554/eLife.42974

Winer JA. 1984. Anatomy of layer IV in cat primary auditory cortex (AI). J Comp Neurol 224:535–567. doi:10.1002/cne.902240405

Winer JA, Miller LM, Lee CC, Schreiner CE. 2005. Auditory thalamocortical transformation: Structure and function. Trends Neurosci. doi:10.1016/j.tins.2005.03.009

Xue Y-X, Xue L-F, Liu J-F, He J, Deng J-H, Sun S-C, Han H-B, Luo Y-X, Xu L-Z, Wu P, Lu L. 2014. Depletion of perineuronal nets in the amygdala to enhance the erasure of drug memories. J Neurosci 34:6647–58. doi:10.1523/JNEUROSCI.5390-13.2014

Yang J-W, Hanganu-Opatz IL, Sun J-J, Luhmann HJ. 2009. Three Patterns of Oscillatory Activity Differentially Synchronize Developing Neocortical Networks In Vivo. J Neurosci 29:9011–9025. doi:10.1523/JNEUROSCI.5646-08.2009

